# Feedback between filament spacing, crosslinker binding, and self-organization in cytoskeletal bundles

**DOI:** 10.64898/2026.07.14.738536

**Authors:** Daniel Steckhahn, Meredith D. Betterton

## Abstract

The lateral spacing between filaments in crosslinked cytoskeletal bundles is a critical yet poorly understood physical parameter that affects force generation, transport, and bundle architecture. We develop a biophysical model of crosslinkers and crosslinking motors on filament pairs. Motor/crosslinker binding sets the filament spacing, which in turn biases which motors/crosslinkers can bind. Crosslinking motors generate pulling forces that bring antiparallel filaments closer together, while non-motor crosslinkers with a preferred binding angle exert repulsive torques that maintain larger spacing. We demonstrate these principles in a model of the fission yeast anaphase mitotic spindle midzone, where microtubules form a square array with nearest-neighbor spacing 2-5 times smaller than the length of crosslinking proteins. Our model shows that motor-crosslinker interactions alone are sufficient to drive self-organization into this experimentally observed geometry. Furthermore, the feedback between geometry and binding creates strong indirect cooperativity, because crosslinkers establish spacing that favors binding of similar-length proteins, leading to history-dependent states that persist long after individual protein binding equilibration. This feedback mechanism in which crosslinkers control geometry and geometry controls crosslinker binding should operate in any multi-crosslinker-filament system and represents a general self-organizing principle for cytoskeletal networks.

## I. INTRODUCTION

Cytoskeletal filament networks generate force and provide structural support important for cell shape, movement, and division [1]. Outside of cells, they are the components of cytoskeletal active matter [2]. Filaments in the cytoskeleton are connected and organized by motor and passive crosslinking proteins. While the role of motors and crosslinkers in generating the architecture of cytoskeletal assemblies has been examined for years, a relatively understudied aspect of cytoskeletal biophysics is the role of the lateral spacing between filaments in force generation, motor and crosslinker binding, and assembly architecture. The force exerted by motors and crosslinkers alters the spacing, organizing filament bundles. Crosslinking protein binding also depends on the spacing between filaments and is decreased if the spacing is large or small relative to the protein length (fig. 1A)[3–5]. While it is known that crosslinking proteins organize filaments in the cytoskeleton, how different proteins collectively establish and regulate spacing remains an open question.

**FIG. 1.**
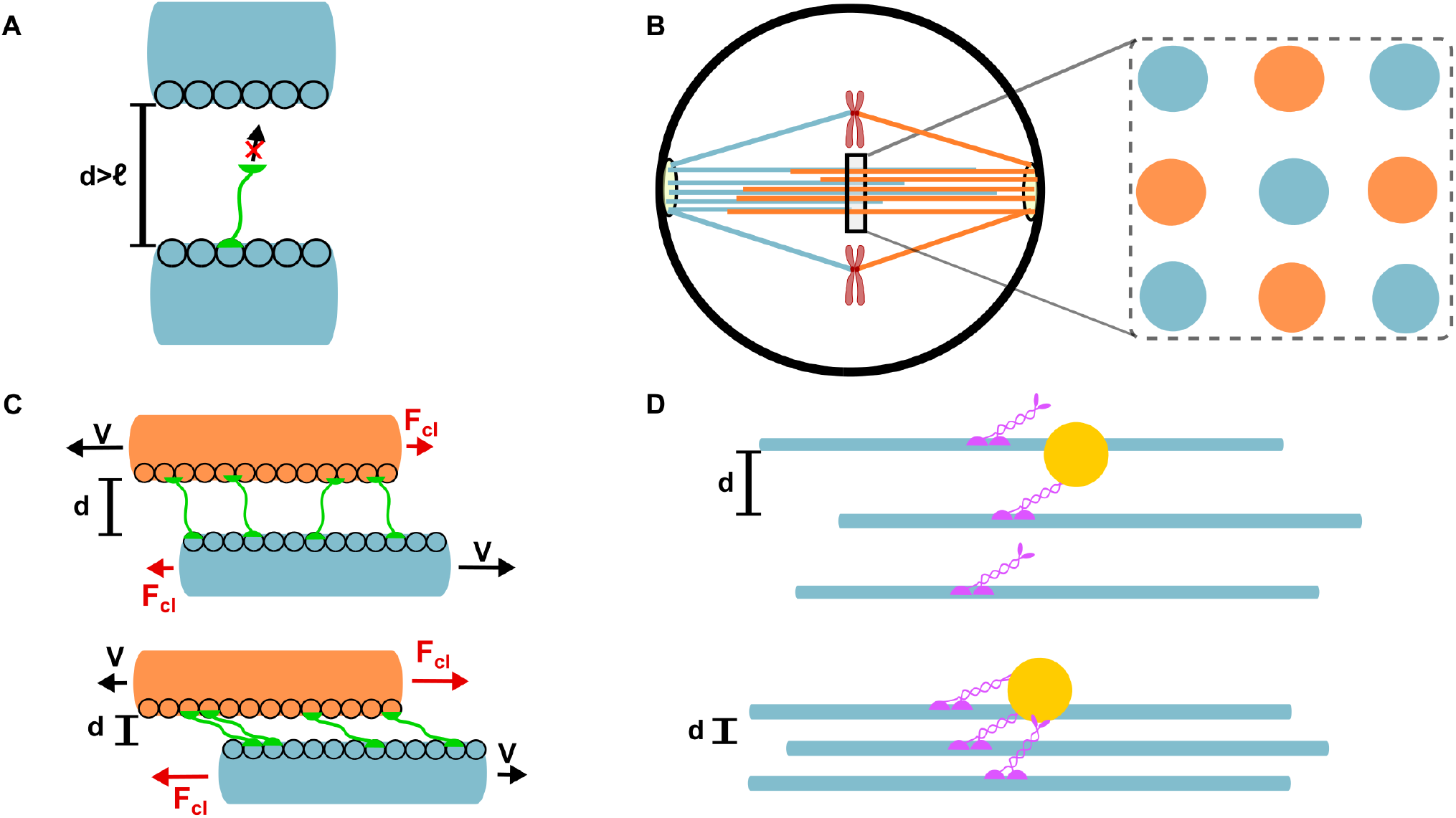
Importance of cytoskeletal filament lateral spacing. (A) Schematic of protein crosslinking inhibition for microtubule spacing greater than the length of the crosslinker. (B) Schematic of the *S. pombe* late-anaphase mitotic spindle and microtubule packing at the midzone.[7, 8] (C) Schematic of resistance to microtubule sliding due to crosslinkers. The resistive force increased as the spacing between the microtubules decreased, dramatically reducing the sliding speed of the microtubules. [13] (D) Schematic of kinesin motor binding and its dependence on microtubule spacing. If the microtubules are close together, multiple kinesins can bind to the same vesicle, increasing the cargo velocity and run length.[14]

In the microtubule-based mitotic spindle, the lateral spacing between microtubules was first measured by electron microscopy beginning in the 1970s. Crosslinked antiparallel microtubules near the center of the spindle typically show a well-defined surface-to-surface spacing, with mean values ranging from 15 nm for the diatom and fission-yeast late-anaphase spindles [6–8], 20 nm in the budding yeast anaphase spindle [9], 25 nm in PtK1 cells [10], and 30 nm in *C. elegans* [11]. Parallel or kinetochore spindle microtubules have a mean 30-nm spacing in the fission-yeast late-anaphase spindle midzone [7, 8], but in other organisms can show a broader distribution or no well-defined spacing [6, 10, 11]. Recent work on human metaphase spindles found that microtubules form locally antiparallel bundles at 8-nm spacing, with similar values for antiparallel and parallel microtubules [12]. What sets the spacing, why it varies between organisms, and the importance for spindle mechanics and integrity [8] are unclear. In the fission yeast *Schizosaccharomyces pombe* mitotic spindle midzone in anaphase, microtubules organize in a square array in which nearest neighbors originate from opposite poles (fig. 1B)[7, 8]. The mechanisms that produce the observed square array remain unexplained.

Lateral spacing has been studied in other contexts in cells. In cilia, changes in the spacing between doublet microtubules have been proposed to coordinate dynein motors during beating [15, 16]. Microtubule bundles in pillar cells from the inner ear have a 42-nm spacing in a square array, structured by crosslinking between microtubules and interdigitated actin [17]. In muscle sarcomeres, the spacing between actin and myosin filaments determines calcium sensitivity and force generation [18, 19], and sarcomere width oscillates with length during spontaneous oscillatory contraction [20].

In reconstituted systems, crosslinker length has been observed to set filament spacing, for example in actin bundles with alpha-actinin [21]. The feedback between spacing and crosslinker binding was shown with actin bundled by two crosslinkers of different length, where the crosslinkers segregate into domains with different spacing [4]. The balance of binding and filament elasticity can explain domain formation [5], showing that the spacing set by one type of crosslinker affects the binding of others. Changing filament spacing also alters myosin II force production in actin bundles [22]. Microtubules crosslinked by the crosslinker PRC1 were observed at spacing close to the crosslinker length [23, 24]. Relative sliding of microtubule pairs crosslinked by the kinesin-5 motor KIF11 showed decreased spacing as motor velocity increased [25]. This is consistent with the intuitive physics of sliding motors, whose force generation to slide filaments depends on their tilt relative to the filament pair. However, for the kinesin-14 motor Ncd spacing did not vary with velocity [26]. Recent work altered crosslinking kinesin-5 motor length with a shortened construct (from 80 to 38 nm long) and found changes in microtubule crosslinking and sliding velocity, consistent with a stiffer crosslinker enforcing a tighter preferred geometry[27]. In microtubule bundles with kinesin-14 motors and depletion interactions, the spacing and lattice organization varied with the level of depletant [28]. The run length of kinesin-1 motors increased along microtubule bundles when crosslinkers were added to increase the spacing of tightly crowded microtubules [29].

Previous theory and computational modeling has addressed aspects of filament spacing in the cytoskeleton, for example to explain segregation of crosslinkers of different length [5]. Modeling of microtubules crosslinked by PRC1 showed spacing resembling experimental measurements [23, 30]. During sliding of crosslinked microtubules, a transition to a low-spacing state dramatically increases the ability of PRC1 to resist sliding apart of microtubule pairs [13, 31] (fig. 1C). Modeling of microtubules in axons found that close interfilament spacing enhances vesicle transport by motors (fig. 1D)[14]. Both the experimental and theoretical work suggest that spacing modulates the mechanical function of crosslinked filaments. However, prior modeling has only begun to address how multiple crosslinking proteins collectively determine spacing, the feedback between spacing and binding, and how these effects organize larger cytoskeletal assemblies.

In this paper, we study physical principles governing lateral spacing in crosslinked filament bundles, and apply them to the *S. pombe* spindle midzone spacing and organization. Using the Coarse-Grained Living Active System Simulator (C-GLASS) [32–35], we model crosslinking proteins on filament pairs, building on prior work on filament and motor/crosslinker modeling [5, 36–47]. We show that non-motor crosslinkers with a preferred binding angle maintain filament spacing close to their rest length. This creates feedback between geometry and binding, because crosslinkers establish spacing that favors binding of similar-length proteins, leading to strong indirect cooperativity and history-dependent states that persist long after individual protein equilibration. Crosslinking motors generate pulling forces that typically bring antiparallel filaments closer together, although for particular parameter ranges large fluctuations in the separation can occur. The force balance between motors and crosslinkers sets the mean separation. Motors and crosslinkers representing the proteins present at the *S. pombe* midzone crosslink microtubules at spacing that closely resembles those found experimentally [7, 8]. Using the lateral forces determined from simulations of microtubule pairs, modeling the full number of microtubules found at the *S. pombe* midzone shows that motor and crosslinker forces together create a square array as seen experimentally [7, 8]. Our results show that the interplay between filament separation and crosslinking protein binding creates a general self-organizing mechanism in cytoskeletal bundles.

## II. MODEL

We developed a biophysical model of cytoskeletal filament pairs and assemblies crosslinked by motors and passive crosslinkers using C-GLASS [32, 35]. The model combines Brownian dynamics for filament motion with kinetic Monte Carlo for stochastic events including crosslinking protein binding/unbinding and stepping/hopping. Because we are primarily modeling microtubules, we assume rigid filaments represented by a single protofilament on the inner edge along which proteins can bind (Fig. 2). Microtubules repel each other through a short-range steric potential and undergo overdamped Brownian motion under forces from motors and crosslinkers (for details, see Appendix A).

**FIG. 2.**
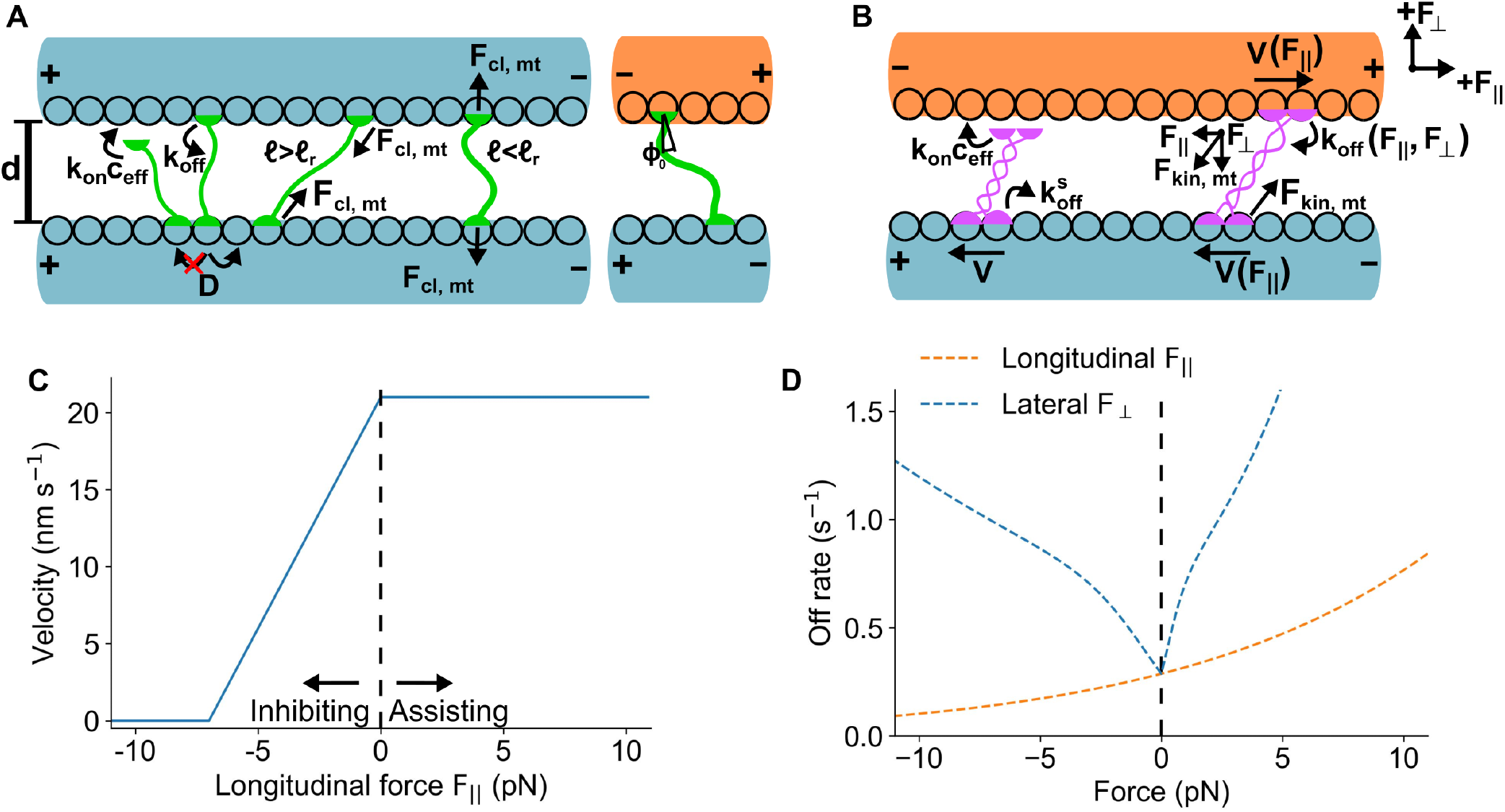
Overview of biophysical model of crosslinking proteins and microtubules. (A) Schematic of crosslinker model. (B) Schematic of crosslinking motor model. (C) Motor force-velocity relation[48, 49]. (D) Motor force-off rate relation[48, 50].

Proteins can bind to one microtubule or crosslink two microtubules (Fig. 2A-B). Crosslinking proteins in solution are represented by a concentration which decreases as proteins bind to the microtubules (Appendix A). Steric interactions prevent bound heads from occupying the same site or crossing when crosslinking [23, 51, 52]. Proteins with one head bound can hop between sites, unbind, or crosslink by binding to a site on the second microtubule. Crosslinking proteins are modeled as two heads connected by a linear spring [13, 30, 38, 53–56]. Each head exerts a force F = −k_s_(*ℓ*−*ℓ*_*r*_) along the tether (molecule axis) between the 2 heads, where k_s_ is the spring constant, *ℓ* the head-to-head distance, and *ℓ*_*r*_ the rest length, giving a spring energy 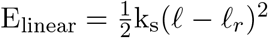. Some crosslinkers, such as the PRC1/Ase1/MAP65 family important in the mitotic spindle show a preferential tilt relative to microtubules [13, 23, 57]. To represent this we model crosslinkers on antiparallel microtubules as torsional springs with energy 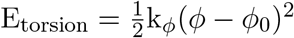, where k_*ϕ*_ is the torsional spring constant, *ϕ* the angle with respect to the microtubule normal, and *ϕ*_0_ the preferred angle. The total crosslinker energy is U = E_linear_ + E_torsion_.

A singly bound protein crosslinks to a site on the second microtubule at a rate 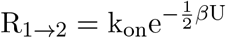, where *β* = 1*/*k_B_T and *U* is the crosslinking energy for binding to that site. This biases crosslinking toward configurations near the spring rest length (Fig. 2A). Crosslinker diffusion, binding, and unbinding transitions obey Boltzmann statistics (Appendix A).

Motors step toward microtubule plus ends (Fig. 2B), with velocity dependent on the longitudinal force on each head through the stall-force relation [48],

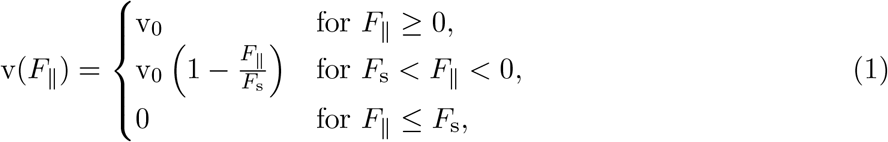

where *F*_s_ is the stall force, *F*_∥_ is the signed force along the walking direction (negative indicates hindering force), and v_0_ is the zero-force velocity (Fig. 2C). For crosslinking motors on antiparallel microtubules, the two heads walk in opposite directions, and when the microtubules slide the heads move relative to each other until the difference in their walking velocities matches the sliding velocity. The longitudinal force at this point is the effective stall force 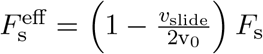, where *v*_slide_ is the relative sliding velocity of the microtubules (Appendix A). Faster sliding lowers the effective stall force. Motor binding rates are force dependent, as for non-motor crosslinkers. Motor unbinding depends asymmetrically on force following kinesin slip-bond behavior under lateral load and catch-bond behavior under longitudinal load [50] (Fig. 2D). We estimated and fit the force-dependent off rate for kinesin-5 from previous work [48, 50] (Appendix A, Fig. S1) [58]. The same unbinding relation was used for kinesin-6, since individual kinesin-6 force-velocity data are not available. Parameter values are given in Table I.

**TABLE I.**
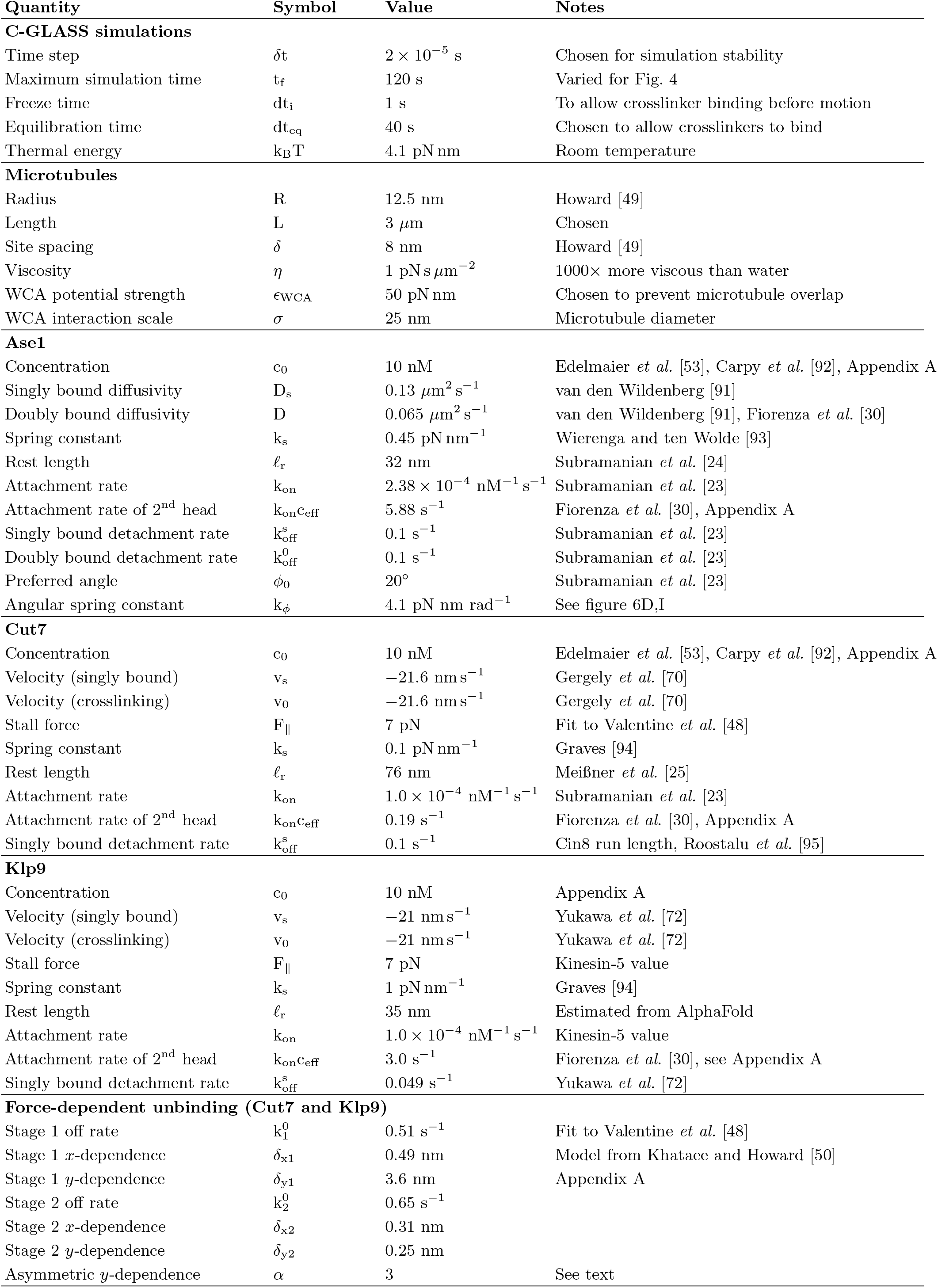
Simulation parameters.

## III. RESULTS

### A. Crosslinkers set microtubule spacing close to their length

We first modeled crosslinkers alone, using parameters based on Ase1/PRC1 [59–61]. For overlaps with short microtubules and a small number of crosslinkers, the free energy as a function of spacing can be computed by explicit enumeration of all possible crosslinking states. Therefore, to compare to semi-analytic calculations, we studied short, 13-site microtubules crosslinked by 5 Ase1 molecules that cannot bind or unbind (Fig. 3A). The model of parallel filaments lacks the torsional energy term, since the crosslinker’s two ends connect at different angles when crosslinking parallel microtubules. For antiparallel microtubules the torsional energy is included. We determined every possible arrangement of one crosslinker bound to an overlap, then every way a second crosslinker could be added without violating steric exclusion, and so on until the desired number of crosslinkers is reached (5 in our example). For every possible arrangement, we calculated the free energy as a function of spacing. The partition function is 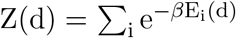, where the sums are over crosslinker configurations. The Helmholtz free energy is H(d) = −k_B_T ln[Z(d)] + U_WCA_(d),where U_WCA_ is the Weeks-Chandler-Andersen potential [62] that models steric repulsion between microtubules. The spacing probability distribution 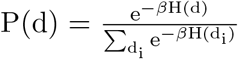, and average spacing 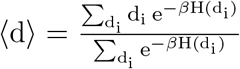, where the sums are over spacing *d*. The force between microtubules 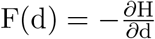.

**FIG. 3.**
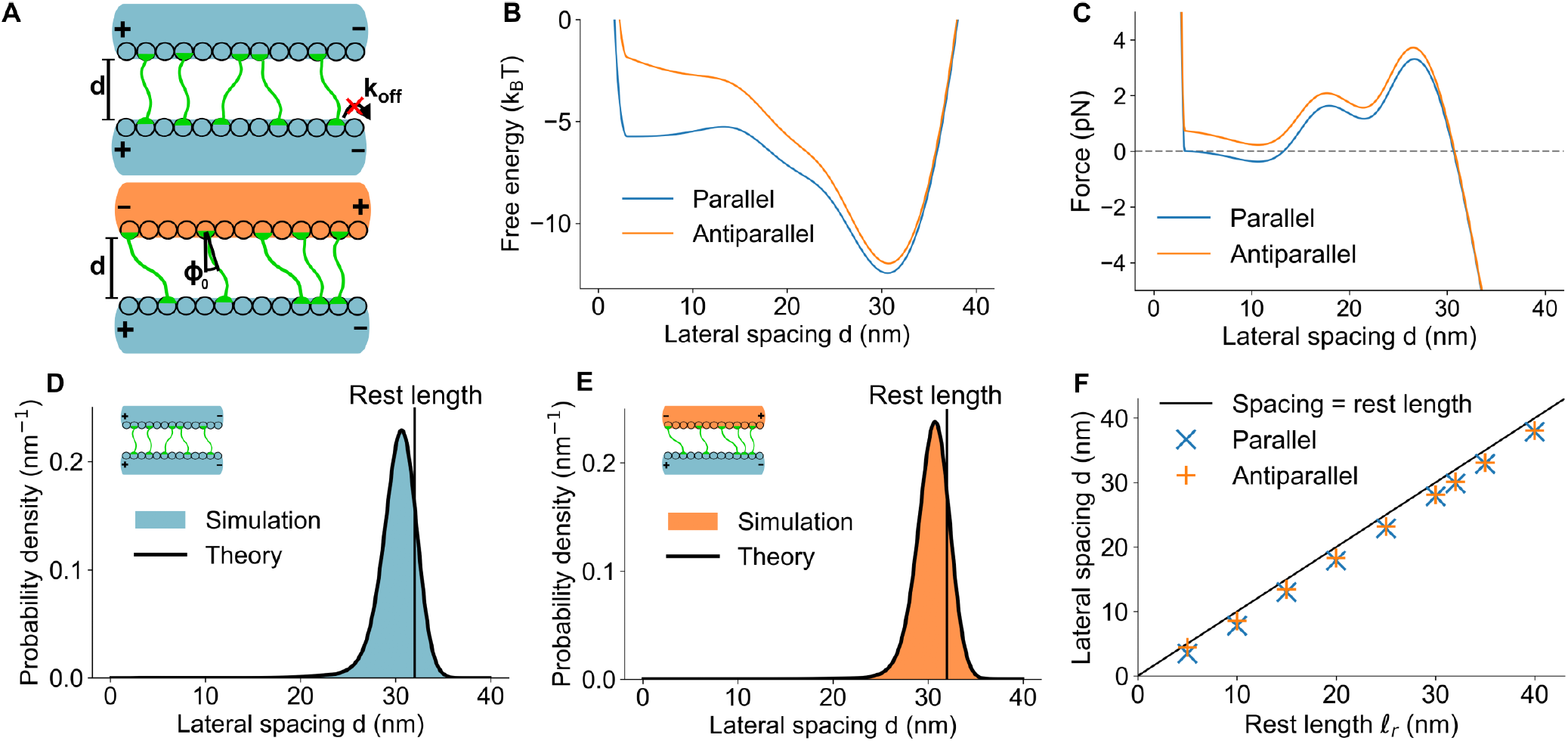
Passive crosslinkers set microtubule spacing near the crosslinker length. (A) Schematic of the model with fixed crosslinker number used for semi-analytic calculation in (B-E). (B) Free energy as a function of spacing for parallel and antiparallel overlaps. (C) Force as a function of spacing for parallel and antiparallel overlaps. (D, E) Spacing probability distribution for parallel (D) and antiparallel (E) overlaps. Colored region, simulation. Black line, semi-analytic theory. Vertical line, crosslinker length. (F) Microtubule spacing as a function of crosslinker rest length on 3 *μ*m overlaps in the model with full binding kinetics. Solid line, spacing equal to the rest length.

The semi-analytical and simulated spacing distributions, free energy, and lateral forces agreed closely (Fig. 3B-E). The average spacing was ≈ 30 nm for both parallel and antiparallel microtubules, slightly less than the 32-nm crosslinker Ase1 rest length [24]. This decrease occurs because for spacing above the rest length, every crosslinker is stretched, pulling microtubules together. At spacing slightly below the rest length, crosslinkers normal to the microtubules are compressed while tilted crosslinkers are stretched, leading to force balance. The distributions of spacing are similar for parallel and antiparallel overlaps, but the force near 15 nm differs: it is repulsive for antiparallel overlaps and at an unstable potential maximum for parallel microtubules. To compare to the full model, we simulated 3-*μ*m-long microtubules with full binding kinetics. The spacing was consistently slightly less than the crosslinker rest length (Fig. 3F). These results suggest that crosslinker force can set the ≈ 30 nm spacing between parallel microtubules at the late anaphase *S. pombe* spindle midzone (Fig. 1), but alone cannot explain the ≈15 nm antiparallel spacing [7, 8].

#### Feedback between microtubule spacing and crosslinker binding leads to history dependence and cooperativity

In systems with actin and two different length crosslinkers, the binding length preference along with actin flexibility led to segregation of crosslinkers into different domains with different filament spacing [4, 5]. To study how this competition between crosslinkers of different length affects binding and spacing on more rigid microtubules, we modeled long (32 nm) and half-length (16 nm) crosslinkers (Fig. 4A). Due to the spring-like crosslinker energy, the rate of crosslinking to a second filament peaks for spacing near the rest length (Fig. 4B,C).

**FIG. 4.**
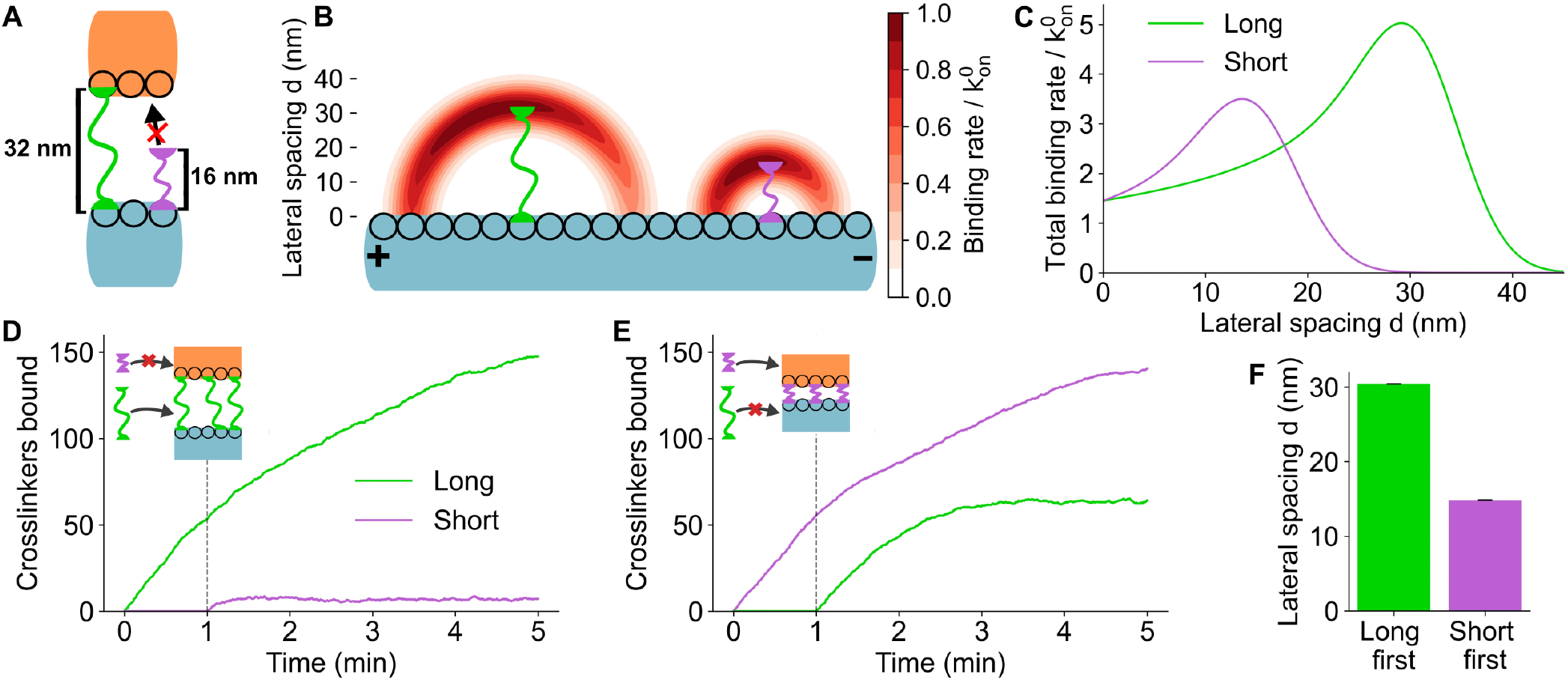
Microtubule spacing affects crosslinker binding, leading to history dependence. (A) Schematic of long and short crosslinkers. (B) The binding rate distribution for long and short crosslinkers in two dimensions, normalized by the bare on rate. (C) The total binding rate of long and short crosslinkers to a second microtubule as a function of spacing, normalized by the bare on rate. (D) The number of long and short crosslinkers bound to a microtubule overlap over time when long crosslinkers are allowed to bind one minute before short. (E) The number of long and short crosslinkers bound to a microtubule overlap over time when short crosslinkers are allowed to bind one minute before long. (F) Average spacing between crosslinked microtubules in the simulations shown in D and E.

Crosslinker binding thus establishes geometry that favors binding of similar-length crosslinkers. To test how this affects systems of mixed crosslinker length, we simulated adding crosslinkers of one length first, followed by the other length later (Fig. 4D,E). The crosslinker added first set overlap spacing, and as a result remained bound at higher number long after the second species was added (Fig. 4D-F). The behavior of crosslinked overlaps therefore depends on the order that proteins are added. This causes history dependence of overlap geometry, and binding inhibition of crosslinkers of different length.

#### Motor proteins pull antiparallel microtubules together

We next modeled microtubules crosslinked by only motor proteins. On antiparallel microtubules, the two heads of a crosslinking motor walk in opposite directions, stretching or compressing the motor and generating both longitudinal force (which slides microtubules) and lateral force (which changes spacing, Fig. 5A). When the spacing is smaller than the motor rest length, motors preferentially bind with a tilt so that their contour length is near the rest length.

**FIG. 5.**
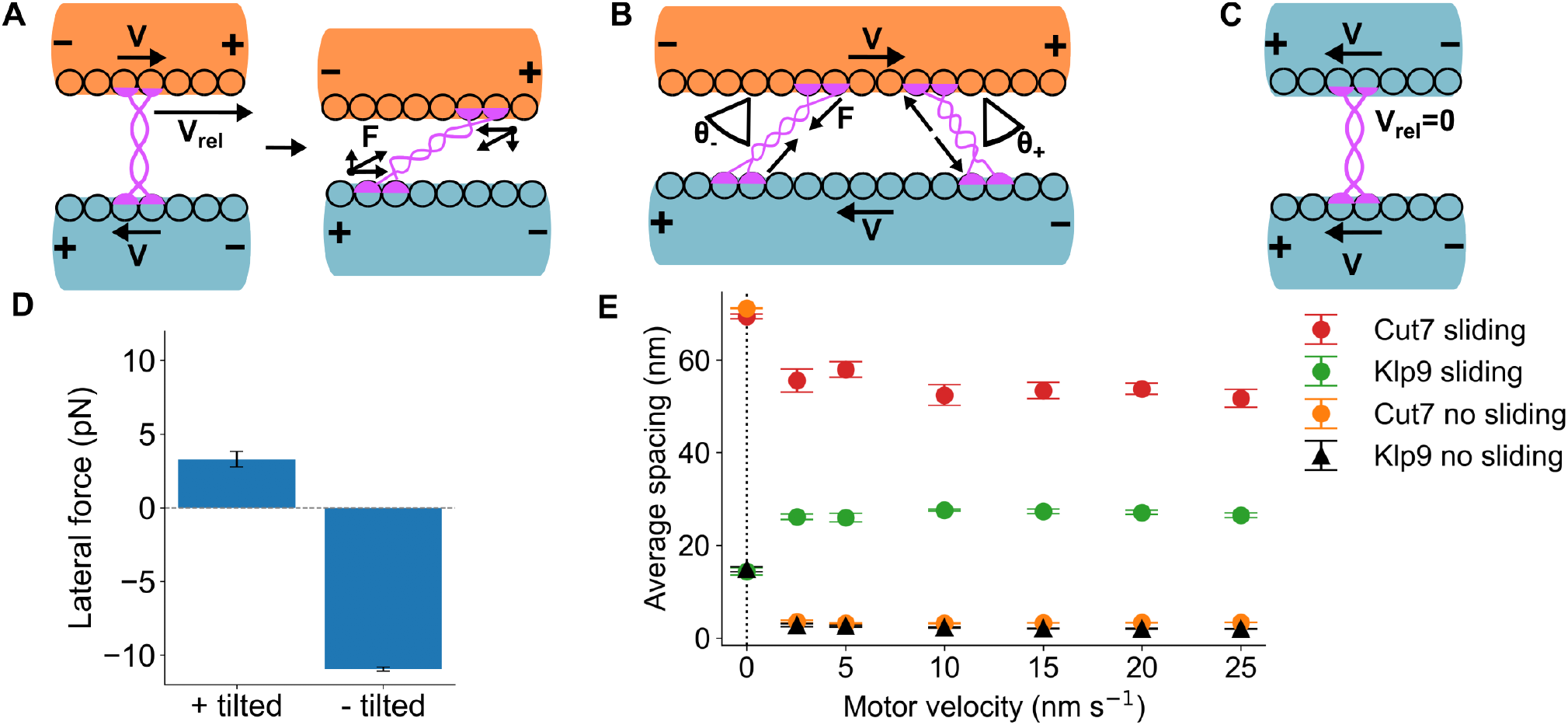
Crosslinking motor proteins pull antiparallel microtubules together. (A) Schematic of force generated by motors crosslinking antiparallel microtubules. (B) Schematic of pulling and pushing force by negatively tilted (left) and positively tilted (right) motors on antiparallel microtubules. (C) Schematic of motor motion on parallel microtubules. (D) Lateral force produced by positively tilted and negatively tilted motors as shown in B. Force was measured in simulations with Klp9 motors at 20 nm/s speed. (E) Microtubule spacing as a function of motor speed for overlaps that were fixed (no longitudinal sliding) and free to slide. The spacing for parallel microtubules is shown at *v* = 0.

The lateral force exerted by a single motor depends on the binding orientation. Motors denoted negatively tilted have a motor tether tilted relative to the head toward the minus end of the filament. For this orientation, stepping tends to stretch the motor tether, driving antiparallel sliding and also producing lateral force that pulls the microtubules together. Negative tilt is typically favored for plus-end-directed sliding motors. Motors denoted positively tilted have the tether tilted relative to the head toward the plus end. In this case, motor stepping compresses the tether and produces lateral force that pushes microtubules apart (Fig. 5B). The longitudinal force is in the same direction for both orientations, sliding the microtubules and reducing the relative head velocity. On parallel microtubules there is no relative head velocity, and sliding does not occur (Fig. 5C).

We modeled the kinesin-6 Klp9 and the kinesin-5 Cut7, which are important for anaphase spindle elongation in fission yeast and localize to the midzone [34, 53, 56, 60, 63–75]. For simulations without sliding, both microtubules were fixed longitudinally, while for sliding simulations one microtubule was allowed to slide (Fig. 5D, E). In the sliding condition, the shorter (3 *μ*m) microtubule began with its minus end adjacent to the plus end of the longer (5 *μ*m) microtubule, resembling the experimental setup of previous work [25]. We varied the motor stepping velocity and kept run length constant as velocity varied by adjusting the motor off rate [76–79]. The asymmetric lateral-force-dependent unbinding model (Fig. 2D, Fig. S1, Appendix A) was used throughout; without it, microtubules fluctuate between large and small spacing (Appendix A, Fig. S2).

As expected, in our model positively tilted motors push microtubules apart while negatively tilted motors pull them together (Fig. 5D). For our model of kinesin-5/Cut7 with sliding, parallel microtubules had larger spacing than antiparallel, and spacing decreased slightly with velocity (Fig. 5E). This velocity-spacing relationship resembled experimental data from the human kinesin-5 KIF11 [25]. On the other hand, in our model of kinesin-6/Klp9 with sliding, parallel microtubules had lower spacing than antiparallel. For antiparallel microtubules with sliding, typically motors produced spacing slightly below the motor rest length. By contrast, non-sliding antiparallel microtubules were driven near zero spacing by either motor, because the higher relative head velocity on longitudinally fixed filaments increased the lateral pulling force. These results show that motors can pull microtubules close together, and that this effect is greatly increased when other forces limit relative microtubule sliding. This effect could contribute to the small spacing of antiparallel microtubules in the mitotic spindle.

#### Force balance between motors and crosslinkers can set intermediate lateral spacing

As a model problem to determine the physical effects that set lateral spacing, we considered the late anaphase spindle midzone in fission yeast. Midzone microtubules form a square array with neighboring antiparallel microtubules separated by ≈ 15 nm and neighboring parallel microtubules by ≈ 32 nm (Fig. 6A) [7, 8]. We modeled the three crosslinking motor and crosslinker proteins important during fission-yeast anaphase B: Klp9, Cut7, and Ase1 [34, 44, 60, 70, 71, 74, 80–82]. Because Klp9 and Cut7 slide antiparallel microtubules and elongate the spindle [74, 82], and sliding affects motor-driven spacing (Fig. 5E), we included sliding in the model (Fig. 6C). One microtubule (6 *μ*m long) was held fixed while the other (3 *μ*m long) slid due to motor and crosslinker forces and a constant opposing external force representing resistance from the spindle pole bodies, chromosomes, nuclear envelope, and other cellular components. When this net external resistive force was 60 pN, microtubules slid at a speed of 16 nm/s, comparable to the measured spindle elongation rate [8, 82, 83].

**FIG. 6.**
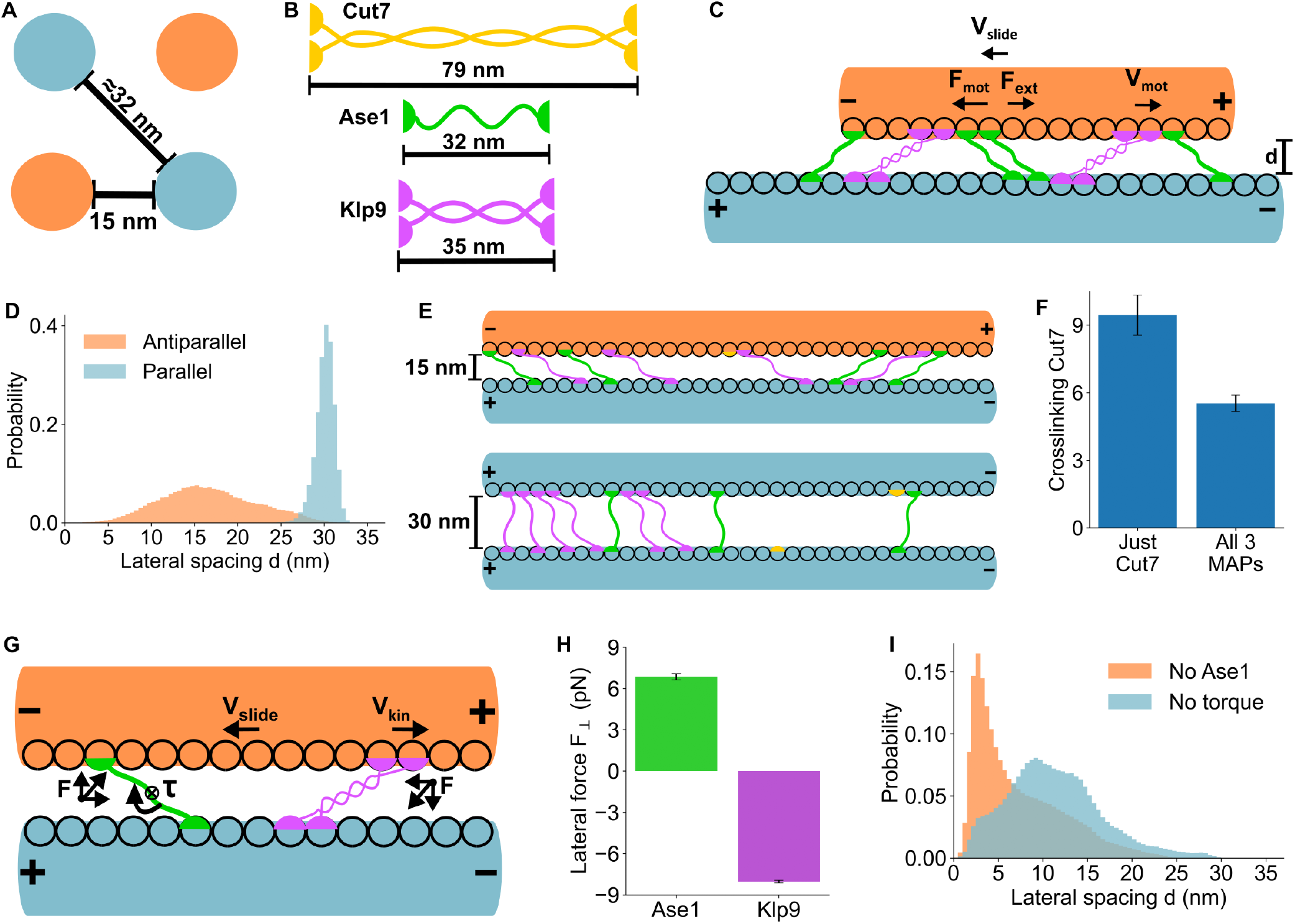
Motors and crosslinker force balance produces spacing similar to that found in the fission-yeast anaphase spindle midzone. (A) Schematic of average microtubule spacing in the *S. pombe* late anaphase spindle midzone[7, 8]. (B) Schematic of motors and crosslinker, with lengths to scale with A. (C) Schematic of simulation geometry. (D) Probability distribution of microtubule spacing for simulations with parallel and antiparallel microtubules. (E) Simulation snapshots showing microtubule, motor, and crosslinker positions from simulations with antiparallel (top) and parallel (bottom) microtubules. (F) The average number of bound Cut7 on overlaps with just Cut7 and with all 3 crosslinking proteins. (G) Schematic showing balance between crosslinker torque that pushes microtubules apart and Klp9 motor force that pulls microtubules together. (H) Lateral force on antiparallel microtubules from the crosslinker Ase1 and motor Klp9. (I) Probability distribution of antiparallel microtubule spacing for simulations lacking Ase1, and with Ase1 modeled without a preferred angle.

With all three crosslinking proteins, microtubule pair spacing in our model was similar to experimental values [7, 8]: 15 nm for antiparallel and 29 nm for parallel microtubules (Fig. 6D, E). Cut7 had difficulty crosslinking because its length would cause it to cross other molecules when binding, which is forbidden in our model. This effect could contribute to the observed decrease in Cut7 midzone localization during anaphase B [34] (Fig. 6E, F). The antiparallel spacing is therefore set by Klp9 pulling microtubules together and Ase1 pushing them apart (Fig. 6G-I). Klp9 pulls constrained microtubules together because the midzone sliding velocity (≈ 16 nm/sec) is closer to fully constrained (0 nm/sec) than to freely sliding Klp9-crosslinked microtubules (≈ 36 nm/sec) (Fig. 5E). Ase1 molecules on antiparallel microtubules are tilted, leading to torsional forces that push microtubules apart (Fig. 6E,G,I, Fig. 3).

Removing Ase1 reduced the average spacing to ≈ 3 nm, confirming that crosslinkers resist motor-driven decrease of spacing. With Ase1 present but lacking an angular preference, the spacing was ≈ 10 nm, showing that torsional forces from tilted crosslinkers help maintain the 15 nm spacing in our model (Fig. 6I). These results are consistent with experimental observations that microtubule spacing in fission-yeast interphase bundles decreased in mutants lacking Ase1 [84]. On parallel microtubules, where crosslinking motors do not pull microtubules together, Ase1 crosslinking maintains spacing close to its rest length, as without motors (Fig. 3). The observed *S. pombe* spindle late-anaphase midzone microtubule spacing can thus be explained by force balance between motors and crosslinkers.

#### Effective pair interactions self-organize an antiparallel square array that closely resembles experimental observations

To test whether motor- and crosslinker-generated forces are sufficient to produce the square array with nearest-neighbor antiparallel microtubules observed in the fission yeast late-anaphase midzone [7, 8], we computed effective pairwise forces between microtubules from our overlap model. We first determined the spacing probability distribution *P* (*d*) using umbrella sampling and the Weighted Histogram Analysis Method (WHAM) to extend the sampled range (Appendix B). We then determined an effective pair interaction free energy *E*(*d*) = −*k*_*B*_*T* ln *P* (*d*) (Fig. 7A) and force by differentiating the free energy *F* (*d*) = −∇*E*(*d*) (Fig. 7B). This is not an equilibrium free energy, because the spacing is determined by nonequilibrium motor activity; instead the effective force represents the tendency of pair interactions with multiple motors and crosslinkers to sample values of the microtubule spacing. We computed this effective force separately for parallel and antiparallel microtubules, using the same conditions as in Fig. 6.

**FIG. 7.**
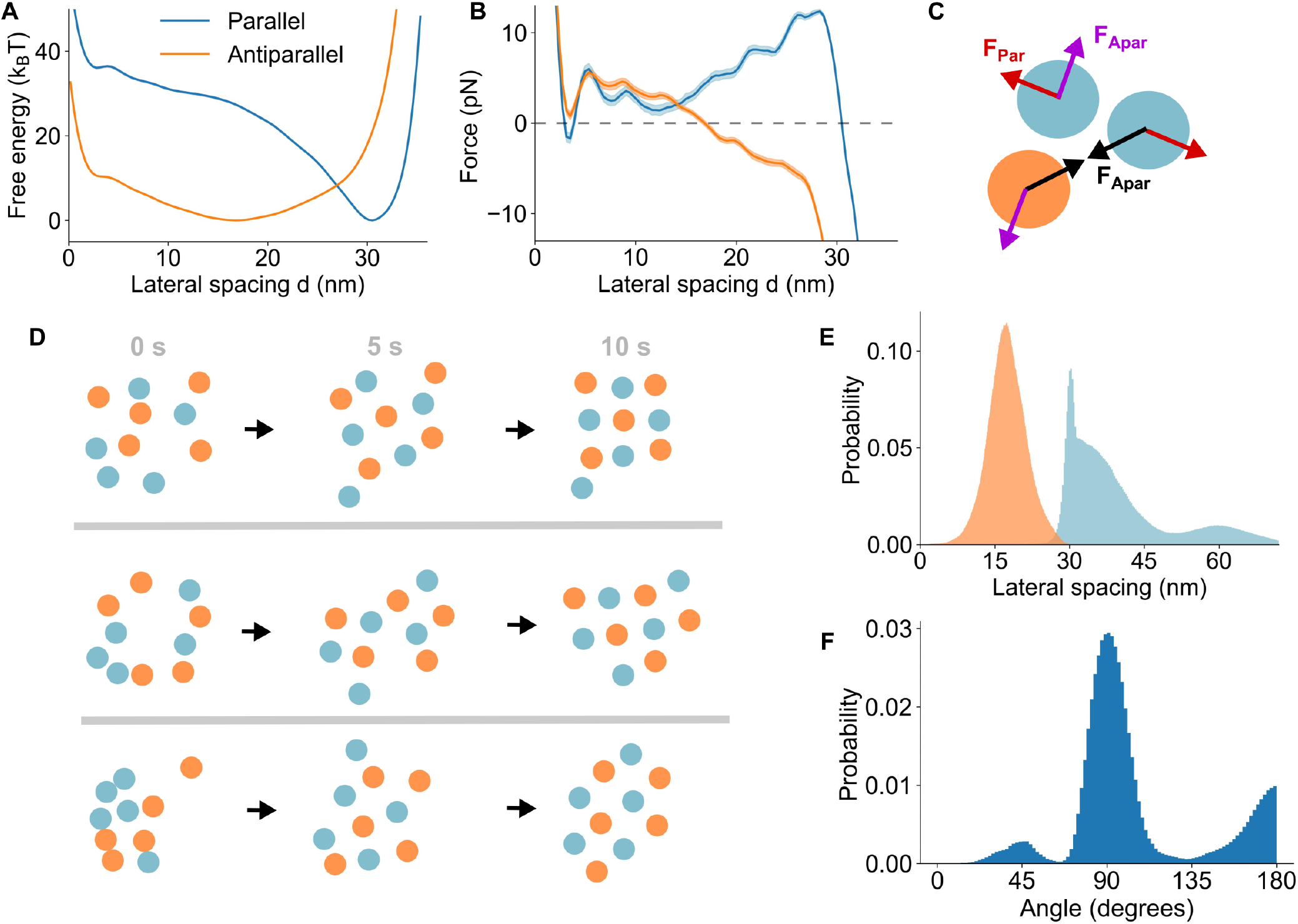
The effective pairwise force between microtubules is sufficient to explain the spacing and organization of the fission-yeast spindle midzone in late anaphase. (A) Effective free energy of crosslinked parallel and antiparallel microtubules as a function of spacing, derived from the spacing probability distributions. (B) The effective lateral force between microtubules as a function of spacing, derived from the free energy shown in A. (C) Schematic of microtubule pair interactions in the multi-filament simulations. (D) Simulation snapshots showing initially randomly positioned microtubules assembling into a square array. The first frame is the initial condition of the simulations, the second after 5 seconds, and the third after 10 seconds of simulation time. (E) The spacing distribution between neighboring parallel and antiparallel microtubules in the multi-filament model. (F) The distribution of the angles between the two nearest neighbors of microtubules in the multi-filament model.

We used this effective pair-interaction force in a 2D model of 10 particles representing microtubules in cross section, with two species (5 from each pole) interacting via the parallel or antiparallel force (Fig. 7C, Appendix B). When initialized with random particle positions, the microtubules self-organized into a square array with antiparallel nearest neighbors (Fig. 7D). The typical spacing was ≈ 15 nm between antiparallel neighbors and ≈ 30 nm between parallel neighbors, matching data from electron microscopy [7, 8] (Fig. 7E). The typical angle between nearest neighbors was 90^°^, as expected for a square array (Fig. 7F). Therefore, the motor and crosslinker interactions that set midzone microtubule pair spacing are sufficient to self-organize the antiparallel square array.

## IV. DISCUSSION

Using a biophysical model of filament pairs with crosslinking proteins, we identified the physical mechanisms that set lateral spacing between pairs. Passive crosslinkers maintain spacing close to their rest length, while crosslinking motors pull antiparallel filaments together, producing spacing well below the motor length. With mixtures of crosslinking proteins, the force balance between motors and crosslinkers sets the steady-state geometry. The spacing in turn controls the accessibility and binding kinetics of crosslinking proteins, creating feedback between filament spacing and crosslinker binding. In the fission-yeast midzone, these pairwise forces are sufficient to self-organize microtubules into the observed antiparallel square array.

The result that passive crosslinkers space filaments near their rest length agrees with measurements in reconstituted systems, in which crosslinker length sets the spacing of actin and microtubule bundles [4, 21, 24]. For future work, a direct test in a motor system would vary the crosslinking motor length. Shortening the kinesin-5 Eg5 from 80 to 38 nm altered microtubule crosslinking and sliding [27], and measuring the pair spacing as a function of motor length would test this prediction.

Because crosslinkers bind most readily when the spacing is near their rest length, the spacing they establish feeds back on crosslinker binding. If two crosslinker species of different length are added to microtubules at different times, we predict the first will remain bound at higher number long after the second is introduced. Shortened crosslinker mutants can be made and their binding measured by fluorescence microscopy [13, 85, 86] in future work. The feedback also produces indirect binding inhibition, because crosslinkers favor spacing near their rest length and promote binding of similar-length proteins. This cooperativity has been observed in crosslinked actin. When fascin and alpha-actinin sort into domains, a mass-action model underpredicts the domain length by a factor of ∼50, and fitting the data required a switching barrier of ∼ 5 *k*_*B*_*T* between the two spacing states [4]. Crosslinker stiffness sets the cooperativity, and above a critical stiffness the bound density undergoes a first-order transition with hysteresis [87]. Our model considered rigid filaments; in flexible helical actin, filament torsion adds a further contribution, so that spacing reflects a balance between crosslinker geometry and filament twist [87]. The overlap stability from this feedback may help cells control which crosslinkers accumulate on cytoskeletal structures and the resulting bundle architecture.

In the mitotic spindle midzone, antiparallel microtubule spacing varies between organisms, from ∼ 8 − 30 nm [6–12]. Our model suggests this variation could arise in part from differences in motor activity. The lateral force with which motors pull antiparallel microtubules together grows when relative sliding is constrained, and the spacing also depends on the motor stepping speed (Fig. 5). The ∼8 nm spacing of human metaphase spindles is well below the ∼80-nm length of a kinesin-5 tetramer and matches the small spacing our model predicts under constrained sliding [12]. In these dense spindles the spacing scaled with microtubule density rather than with the rest length of any single crosslinker [12]. Our mechanism assumes that pairwise motor and crosslinker forces dominate over collective steric and depletion interactions, so this density scaling and the absence of a parallel-antiparallel difference may be due to effects beyond the pair limit. The pair force balance applies most directly in sparser arrays such as the fission-yeast midzone, where the crosslinker and motor population is small and well characterized.

We tested these principles quantitatively in a model of the fission-yeast late-anaphase spindle midzone. Its structure was first measured more than 30 years ago, but the mechanisms that set it had remained unexplained [7, 8]. We find that the square array and parallel and antiparallel spacing can be explained by force balance of midzone crosslinking proteins. The same model predicts spacing for other motor and crosslinker combinations. For example, cells lacking the crosslinker Ase1 in *S. pombe* should have lower microtubule spacing than wild type, consistent with the reduced spacing seen in interphase bundles lacking Ase1 [84].

These principles should operate in any system where crosslinker geometry and binding interact. In muscle sarcomeres, titin and other crosslinkers bridge thick and thin filaments at a spacing that depends on sarcomere length [18, 20]. Changes in this spacing could shift the balance of bound motors and crosslinkers and contribute to length-dependent force generation. In cilia, dynein motors and nexin crosslink adjacent doublet microtubules [15, 16], and the same motor and crosslinker force balance could set the interdoublet spacing that controls beating. In reconstituted active matter, where mixtures of motors and crosslinkers self-organize filament networks [22], spacing-dependent binding could produce history-dependent structures with geometry that depends on the order of protein addition.

In summary, the interplay between filament spacing and crosslinking protein binding is a general self-organizing mechanism in cytoskeletal bundles. The force balance between motors and crosslinkers sets the geometry, and the geometry in turn biases which proteins bind. Because the spacing established by one protein favors binding of similar proteins, filament geometry depends on assembly history and acts as a memory that biases later binding. These pairwise interactions are sufficient to self-organize filaments into the ordered arrays seen in the spindle midzone. These principles apply to crosslinked filament systems throughout the cytoskeleton.

## ACKNOWLEDGMENTS

We thank Richard McIntosh, Matthew Glaser, Shane Fiorenza, Adam Lamson, and Jeffrey M. Moore for helpful discussions. This work was supported by National Science Foundation grant No. DMS-2153399 to M.D.B. This work used the Alpine supercomputer, jointly funded by the University of Colorado Boulder, the University of Colorado Anschutz, Colorado State University, and the National Science Foundation (awards 2201538 and 2322260).

## Appendix A Model and C-GLASS simulations

This appendix provides details of the model and simulation in C-GLASS [32–35]. The code is available at https://github.com/Betterton-Lab/C-GLASS.

### 1. Microtubules

Microtubules are modeled as rigid filaments because in this work they are typically much shorter than the persistence length [88]. One protofilament with tubulin sites spaced 8 nm apart is modeled on the inner edge of each microtubule (Fig. 2). The microtubules undergo Brownian dynamics. During each time step the lateral displacement is

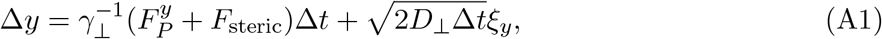

When free to slide, the longitudinal displacement is

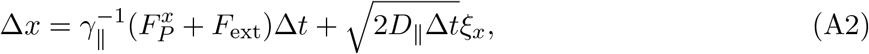

where *ξ*_x_ and *ξ*_y_ are independent standard normal random variables, 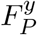 and 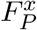 are the lateral and longitudinal components of the force exerted by crosslinking proteins, *F*_steric_ is the steric force between microtubules and *F*_ext_ is the external force if present. The longitudinal and lateral components of microtubule drag are

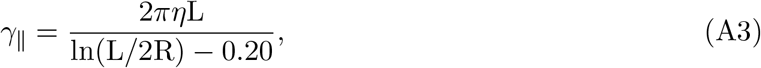

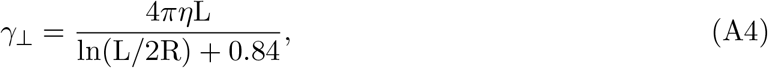

where *η* is the viscosity, L is the length of microtubules, and R is the radius of the microtubule [49, 89]. The longitudinal and lateral components of microtubule diffusion are calculated from the drag using the Einstein relation

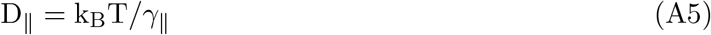

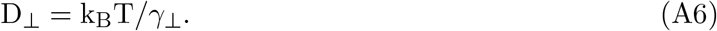

In addition to forces from crosslinking proteins, the microtubules repel each other through a WCA potential representing steric interactions

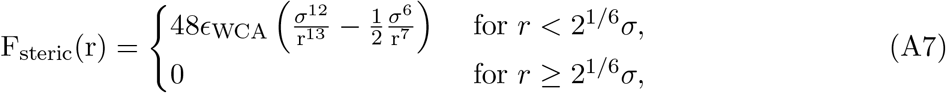

where r is the center-to-center distance between microtubules, *ϵ*_WCA_ is the strength of the potential, and *σ* is the interaction distance.

### 2. Crosslinking proteins

The crosslinker spring force, torsional energy, and crosslinking rate are given in the main text. The torsional spring was only implemented for crosslinkers on antiparallel microtubules. On parallel microtubules the orientation that crosslinkers adopt is unclear, since the two heads connect to microtubules at different angles.

Crosslinking proteins bind from solution to each unoccupied site at a rate 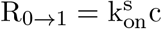, where k_on_ is the binding rate constant and *c* is the protein concentration, which decreases as proteins bind to the overlap as

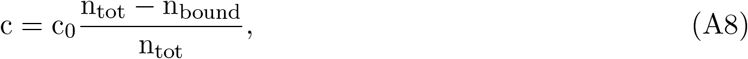

where c_0_ is the initial concentration, n_tot_ is the total number of proteins of a given species, and n_bound_ is the number of bound proteins. The total number of crosslinking proteins of a given species is n_tot_ = c_0_V_cell_*/*20, where V_cell_ is the volume of the cell. The factor of 20 accounts for the ≈ 20 connections between neighboring microtubules at the *S. pombe* midzone; we model one connection at a time [7, 8].

For a crosslinking protein with one head bound, the on rate of the second head crosslinking to the other filament is k_on_ = k_on_c_eff_, where c_eff_ is the effective concentration of a single crosslinking protein within its binding radius. The effective concentration was c_eff_ = 1/V_explored_, where V_explored_ is the volume explored by the unbound head, which was calculated as[3],

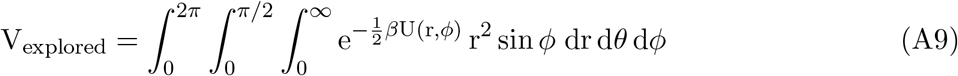

where *ϕ* is the azimuthal angle and *θ* is the polar angle. From the one-head-bound state, crosslinking proteins unbind at a constant rate 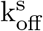. Hopping between sites and unbinding from the crosslinking state are handled differently for motor and non-motor crosslinking proteins as described below.

### 3. Crosslinkers

While bound to a single microtubule, non-motor crosslinking heads diffuse between sites with a hopping rate k_dif_ = D_s_*/δ*^2^, derived from previous work [90] to match the experimentally measured diffusion constant D_s_ [91], where *δ* is the spacing between lattice sites. Because crosslinkers are passive, their diffusion, binding, and unbinding while crosslinking obey Boltzmann statistics.

While crosslinking, the diffusive hopping rate depends on the torsional and spring energy, with 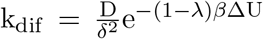, where D is the base diffusion coefficient of a crosslinked head, ΔU is the change in energy between the current and target sites, and *λ* is a constant between 0 and 1 that weights the rates between forward and backward transitions. We set *λ* to 0.5, meaning both forward and reverse hopping are equally weighted. While crosslinking, non-motor crosslinkers unbind at a rate 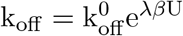.

### 4. Motor proteins

Kinesin motors step toward plus ends at a rate k_+_ = v*/δ*, where *v* is the motor velocity and *δ* is the site spacing. While singly bound, v = v_s_ is constant. When one head is bound, the velocity depends on force through the stall force relation given in the main text.

The force-dependent off rate for kinesin-5 was obtained by dividing published velocity data by run length data point-wise [48], yielding an asymmetric curve similar to what was reported for kinesin-1 (Fig. S1) [50]. This off rate was well fit using the functional form of previous work [50] (Fig. S1). In single-molecule optical trap assays, only forces directed away from the microtubule are measured [50], so it is unclear whether motor unbinding responds symmetrically to positive and negative lateral forces. With a symmetric response, in our model overlaps fluctuated between small (≈ 0) and moderate (≈ motor rest length) spacing (Sec. A 4 b below). Increasing the off rate for positive lateral force relative to negative lateral force by a factor *α* stopped this fluctuation and led to stable spacing.

#### a. Effective motor stall force during filament sliding

As measured experimentally, the stepping velocity of motor protein heads follows the stall force relation

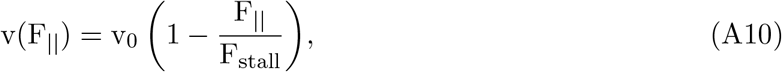

where F_stall_ is the stall force without sliding, F_||_ is the longitudinal force, and v_0_ is the maximum walking speed of motors. Without sliding, motor proteins walk until the longitudinal force, F_||_, reaches the stall force,

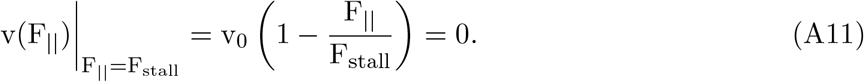

For crosslinking motors on antiparallel microtubules, the total velocity difference between the motor heads is twice this velocity because the heads walk in opposite directions. When filaments are free to slide, the motor heads move relative to each other until the velocity difference from walking is the sliding velocity

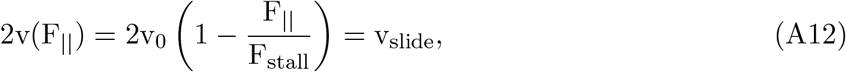

where v_slide_ is the difference in sliding velocity between the two microtubules. Solving for F_||_ gives,

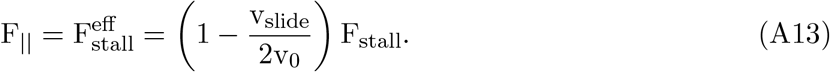

#### (b) Motor-driven fluctuation of spacing

Not all motor parameters lead to stable spacing. If motor unbinding depends symmetrically on lateral force, motors can cause the spacing to fluctuate stochastically (Fig. S2). This occurs because a negatively tilted motor with heads that walk toward the plus end exerts forces that pull the microtubules together. For a positively tilted motor, as the heads walk toward the plus ends the force magnitude (directed against walking) will increase until it reaches a maximum (Fig. S2A). If this maximum force exceeds the modified stall force (Eqn. A13), the motors cannot walk further and remain fixed with positive tilt. Positively tilted motors at the stall force push with greater lateral force than negatively tilted motors at the stall force pull (Fig. S2A) because the positively tilted motors are compressed closer to the microtubule normal, making the lateral component a larger fraction of the total force (Fig. S2B,C). Therefore, for equal populations of stalled positively and negatively tilted motors, the microtubules are pushed apart.

As the spacing increases, the force can fall below the effective stall force and positively tilted motors can walk past normal and become negatively tilted. The increased pulling by negatively tilted motors causes the spacing to decrease until the distance between microtubules drops, in some cases reaching close to zero (Fig. S2D-E). For small spacing, newly bound positively tilted motors will stall before reaching normal. The number of positively tilted motors will then increase until there are enough positively tilted motors to push the microtubules apart, increasing the spacing again. Spacing fluctuations can also occur for motor parameters where motors are unable to walk from positive to negative tilt, due to stochastic fluctuations in the number of positively and negatively tilted motors (Fig. S2F-G).

Spacing fluctuations produce a broad distribution with peaks at large and small spacing and a dip at intermediate spacing (Fig. S2E,G). The consistent spacings found at the *S. pombe* midzone suggest that these large fluctuations are likely not occurring. However, our model suggests that motor-driven spacing fluctuation may occur in other situations or with other motor proteins. The asymmetry in the dependence of motor off rates on lateral force in our default parameters favors negatively tilted crosslinking and prevents this fluctuation.

#### c. Maximum longitudinal force on motors

Motors that are positively tilted encounter a maximum longitudinal force as they walk toward filament plus ends. The longitudinal force is

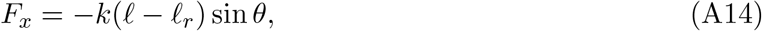

where *k* is the tether spring constant, *ℓ* is the tether length, and *θ* is the angle between the motor tether axis and the normal to the microtubule, signed positive for tilt of the motor tether toward the plus end. Writing *ℓ* in terms of the angle gives

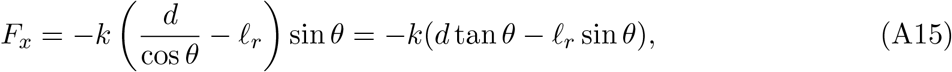

where *d* is the spacing. The force is at an extremum when its derivative is zero,

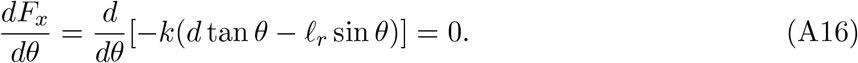

Taking the derivative and simplifying we have

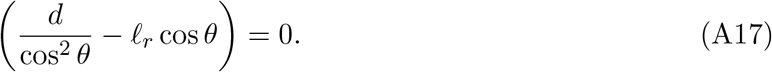

Solving for *θ* gives

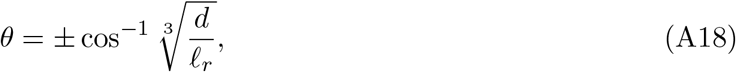

where the positive solution is for positively tilted motors and the negative solution for negatively tilted motors. Plugging the positive root into Eqn. A15 for F_x_ gives

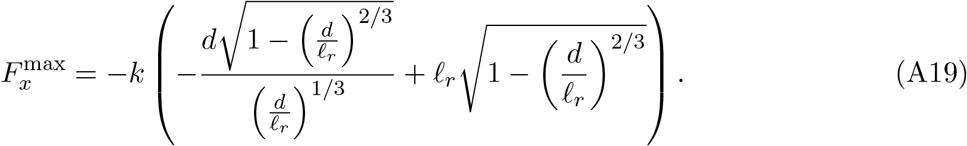

The magnitude of this force is shown in Fig. S2C.

## Appendix B Midzone array model

For simulations of multiple microtubules representing the *S. pombe* late-anaphase spindle midzone, microtubules interact through a force calculated from the effective free energy found in pair simulations (Fig. 7). The free energy was calculated using the spacing probability distribution from the C-GLASS overlap simulations. During the simulations, overlaps explored a narrow range of spacing, resulting in undersampling outside this range. To extend the sampled range, the microtubules interacted through an additional harmonic potential of the form 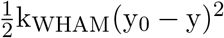, where y_0_, the center of the potential, was varied in steps of 5 nm between 0 and 50 nm, and k_WHAM_, the strength of the potential, was set to 3.29 pN/nm. These simulations were then combined and the free energy found using the weighted histogram analysis method (WHAM) [96].

Once equilibrated, the microtubules were never found between ≈33 nm and ≈47 nm because Ase1 and Klp9 would bind and bring the microtubules closer than 33 nm, or the WHAM potential would hold the spacing high enough that crosslinkers would never consistently bind. Because of this, the free energy did not extend past 33 nm for parallel or antiparallel microtubules. From this free energy, the effective force between microtubules was calculated as 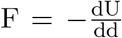, separately for parallel and antiparallel overlaps.

The effective force as a function of microtubule spacing was then used to run simulations with 10 particles representing microtubules, an amount typically found at the midzone of *S. pombe* in late anaphase [7, 8]. There were two species of particles, with 5 particles each, representing microtubules originating from opposite spindle poles. Simulations were run in two dimensions in the plane representing the cross-section of the mitotic spindle. The microtubules were inserted randomly and constrained to move within a 175 × 175 nm box.

Microtubules of different species interacted through the force derived from antiparallel overlaps, and microtubules of the same species interacted through the force derived from parallel overlaps. Microtubules did not interact if a third microtubule was within one microtubule radius *R* of the line connecting their centers. Microtubules diffused and moved in two dimensions using the same *γ*_⊥_ and D_⊥_ as in the C-GLASS simulations for 3 *μ*m microtubules. Simulations were equilibrated for 5 seconds, long enough for the microtubules to arrange into a square array, then data on spacings and angles between microtubules were collected for 15 seconds.

