## Supplemental figures for "Feedback between filament spacing, crosslinker binding, and self-organization in cytoskeletal bundles"

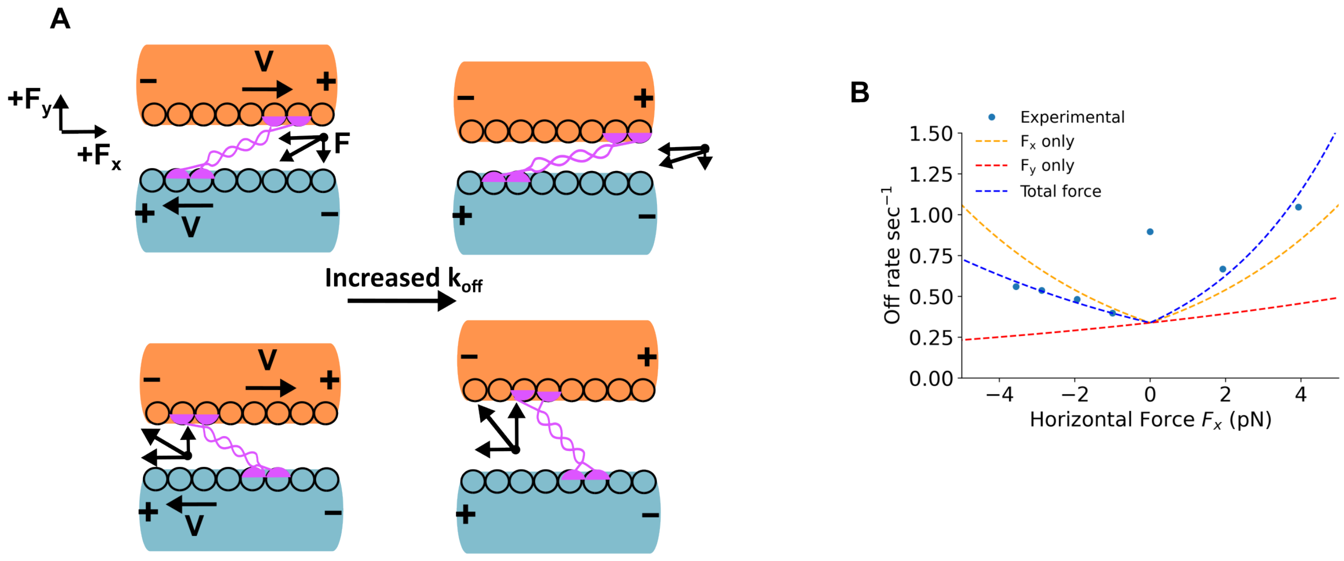


FIG. S1. Effects of lateral force on kinesin binding. (A) Schematic illustrating that the off rate of plus-angled motors increases as spacing decreases, while the off rate of minus-angled motors decreases as spacing decreases. (B) Measured kinesin off rate compared to our binding model. Curves illustrate how the off rate changes due to the longitudinal force, the lateral force, and the total force.

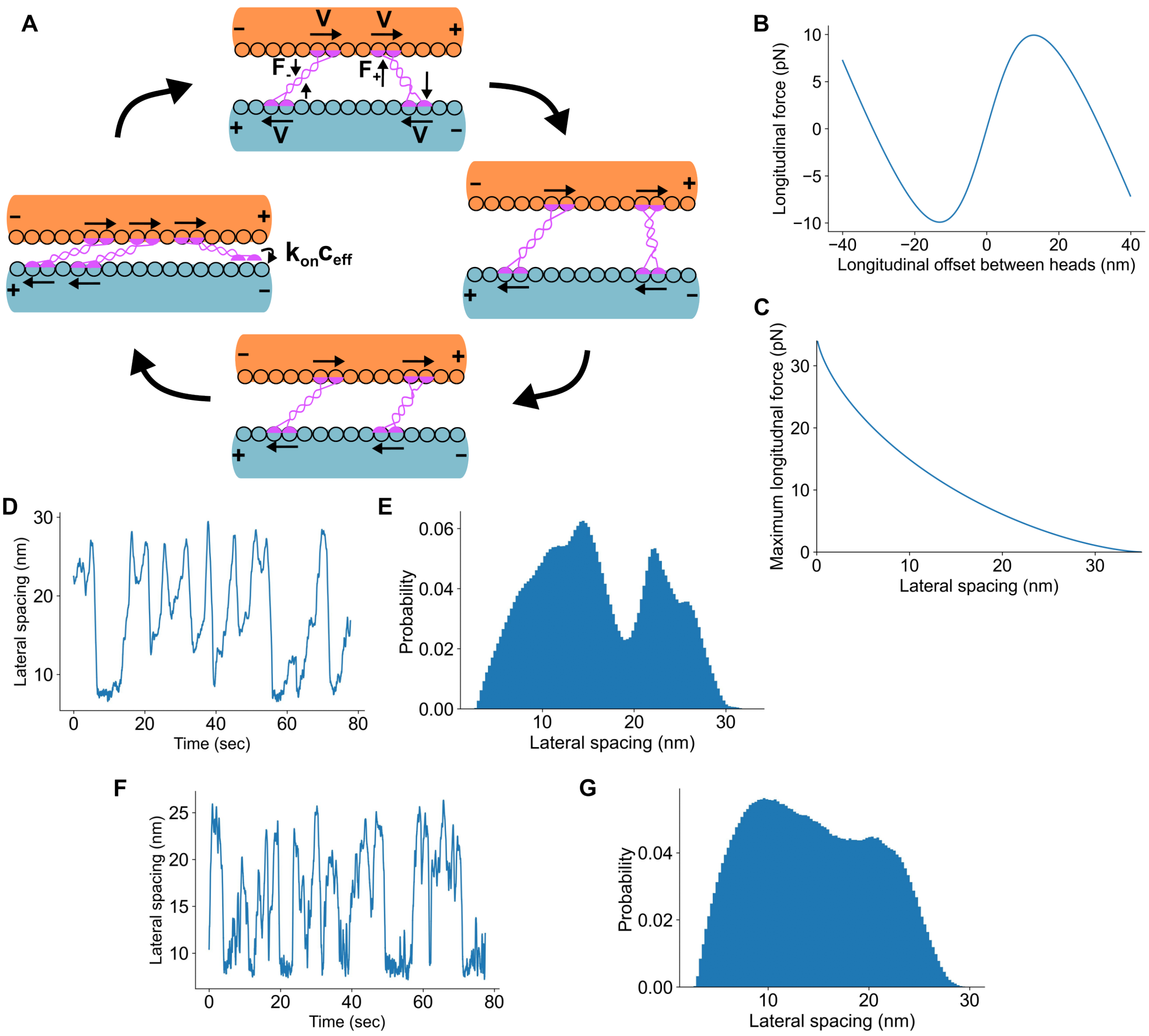


FIG. S2. Motors can drive large fluctuations of spacing. (A) Schematic showing how shifting force balance between positively tilted and negatively tilted motors can cause spacing fluctuations. (B) Longitudinal force on motor heads as a function of the longitudinal offset between the two motor heads at a microtubule spacing of 15 nm. (C) The maximum hindering force on motors walking along microtubules as a function of spacing. If the effective stall force is greater than this value, motors can walk from negatively tilted to positively tilted. (D) Spacing as a function of time from an example simulation with spacing fluctuations caused by motors walking from positive tilt to negative tilt. Simulations used the same setup as the antiparallel simulations in main text Fig. 5 without sliding and with Klp9. Klp9 used the default parameters, except with the stall force halved to 3.5 pN and the off-rate only dependent on longitudinal forces. (E) Spacing distribution during spacing fluctuations for simulations described in D (n=10 runs). (F) Spacing as a function of time from an example simulation with spacing fluctuations with motors unable to step from positive to negative tilt. Simulations used the same setup as the antiparallel simulations in Fig. 6, except with only Klp9. Klp9 used the default parameters, except the lateral force dependence was symmetric ($\alpha = 1$). (G) Spacing distribution during spacing fluctuations for simulations described in F (n=10 runs).
